# Characterization of transgenic mouse lines for selectively targeting glial cells in dorsal root ganglia

**DOI:** 10.1101/2020.02.10.941625

**Authors:** Yasmine Rabah, Bruna Rubino, Elsie Moukarzel, Cendra Agulhon

**Author notes:** These authors contributed equally to this work (Yasmine Rabah, Bruna Rubino, Elsie Moukarzel).

## Abstract

The importance of glial cells in the modulation of neuronal processes is now generally accepted. In particular, enormous progress in our understanding of astrocytes and microglia physiology in the central nervous system (CNS) has been made in recent years, due to the development of genetic and molecular toolkits. However, the roles of satellite glial cells (SGCs) and macrophages – the peripheral counterparts of astrocytes and microglia – remain poorly studied despite their involvement in debilitating conditions, such as pain. Here, we characterized in dorsal root ganglia (DRGs), different genetically-modified mouse lines previously used for studying astrocytes and microglia, with the goal to implement them for investigating DRG SGC and macrophage functions. Although SGCs and astrocytes share some molecular properties, most tested transgenic lines were found to not be suitable for studying selectively a large number of SGCs within DRGs. Nevertheless, we identified and validated two mouse lines: (i) a CreERT2 recombinase-based mouse line allowing transgene expression almost exclusively in SGCs and in the vast majority of SGCs, and (ii) a GFP-expressing line allowing the selective visualization of macrophages. In conclusion, among the tools available for exploring astrocyte functions, a few can be used for studying selectively a great proportion of SGCs. Thus, efforts remain to be made to characterize other available mouse lines as well as to develop, rigorously characterize and validate new molecular tools to investigate the roles of DRG SGCs, but also macrophages, in health and disease.

## Introduction

Astrocytes and microglia serve essential support and immune functions, and contribute to diseases of the CNS (1). For a long time, their heterogeneity and roles in the CNS have remained unclear due to the lack of tools to specifically identify them, and monitor or alter their activity. However, over the last fifteen years, new genetically-encoded tools to selectively visualize astrocytes and microglia as well as read out or abolish astrocytic Ca^2+^ activity have been generated. They include (i) transgenic mice expressing green fluorescent protein (GFP) under the control of astrocytic or microglial promoters (2–5), (ii) transgenic mice expressing genetically-encoded Ca^2+^ indicators (GCaMP) to monitor astrocyte Ca^2+^ dynamics (6–8), or (iii) IP_3_R2 knockout mice to abolish global G_q_ protein-coupled receptor (G_q_ GPCR)-mediated Ca^2+^ elevations in astrocyte cell bodies and large processes (9, 10). These tools have been extremely useful for probing astrocyte and microglia functions in the CNS and are now extensively used in the field.

In the peripheral nervous system (PNS), SGCs in DRGs share several properties with astrocytes, including expression of cytosolic proteins [*e.g.* glial fibrillary acidic protein: GFAP (11); S100 calcium-binding protein beta: S100β (12); glutamine synthetase (13)], membrane neurotransmitter transporters [*e.g.* glutamate aspartate transporter: GLAST (13)], and channels [*e.g*. inwardly rectifying K^+^ channel 4.1 (Kir 4.1) (14); connexin 43 (Cx43)-based gap junctions and hemichannels (15)]. As astrocytes and many other cell types, SGCs use Ca^2+^ as a signaling molecule (16). Additionally, there is substantial evidence for a SGC role in chronic pain wherein SGCs undergo a reactive gliotic response accompanied with increased GFAP expression, hypertrophy, proliferation and upregulation of Cx43 (17, 18). Furthermore, the counterparts of CNS microglia in DRGs are the macrophages, and both express the ionized Ca^2+^-binding adapter molecule 1 (Iba1) (19, 20). Emerging evidence has implicated the contribution of DRG macrophages to neuropathic pain development and axonal repair in the context of nerve injury (21–23). Thus, understanding how SGC and macrophage morphology and function are remodeled in physiology and pathology can help to find new therapeutic targets for pain-related diseases (13,24–26). However, SGC and macrophage heterogeneity and role remain largely unknown in DRGs, mainly due to the unavailability of tools to specifically visualize DRG glial cells and examine or manipulate their Ca^2+^ dynamics. Satellite glial cell or macrophage specific gene expression or deletion would therefore help to clarify the roles of those cells in DRGs.

To accomplish this, using immunohistochemistry or 2-photon Ca^2+^ imaging, we have characterized different available genetically-modified mouse lines widely used to study astrocytes or microglia. Our results showed that most lines used for examining astrocyte functions are very inefficient for studying selectively a large proportion of SGCs in DRGs. However, we have identified two mouse lines allowing either the selective expression of a Ca^2+^ biosensor in SGCs or the labelling of macrophages.

## Material and Methods

### Animals

Mice were housed in transparent plastic cages (5 mice/cage) and fed *ad libitum*. Illumination was controlled automatically with a 12/12h light-dark cycle. S100β-eGFP (5), ALDH1L1-eGFP (27) and CX3CR1-eGFP (3) transgenic lines were used. Furthermore, CAG-lox-STOP-lox-GCaMP6f transgenic mouse line (7) was crossed with GLAST-CreERT2 (28), GFAP-Cre (29), Cx30-CreERT2 (28) or Cx43-CreERT2 (30) mice. As a result, we obtained four new double transgenic mouse lines that we named GLAST-CreERT2::GCaMP6f, GFAP-Cre::GCaMP6f, Cx30-CreERT2::GCaMP6f and Cx43-CreERT2::GCaMP6f. To induce expression of the Ca^2+^ biosensor GCaMP6f, GLAST-CreERT2::GCaMP6f and Cx30-CreERT2::GCaMP6 mice were injected intraperitoneally (i.p.) with tamoxifen (1 mg/day, Sigma) diluted in corn oil (Sigma) during 5 consecutive days. To obtain optimum GCaMP6f expression in brain astrocytes of Cx43-CreERT2::GCaMP6f mice, tamoxifen treatment lasted 10 days as previously described (31). Animals were then used 2 weeks after the first day of treatment. All lines used were kept heterozygous for transgenes encoding GFP/eGFP, GCaMP6f, Cre or CreERT2. Experiments were conducted in 2 to 3 month-old male and female mice from the C57BL/6JRj background. Animal care and procedures were carried out according to the guidelines set out in the European Community Council Directives. The protocol was approved by the Committee on the Ethics of Animal Experiments of Paris Descartes University (Protocol Number: 2018061412588713).

### Immunohistochemistry, image acquisition and analysis

Animals were transcardially perfused with 4% paraformaldehyde under ketamine/xylazine (100 mg/kg – 10 mg/kg respectively, i.p.) anesthesia. Brains and lumbar L3, L4 and L5 DRGs were removed, post-fixed for 24 h or 2 h in 4% paraformaldehyde, respectively. Then, tissues were cryoprotected overnight at 4°C in 0.02 M phosphate buffer saline (PBS, pH 7.4) containing 20% sucrose, and frozen in optimal cutting temperature compound. Sixteen or 14 µm thick sections (brain or DRG, respectively) were cut using a cryostat (Leica), mounted on Superfrost glass slides and stored at −80°C. The day of the experiment, sections were washed 3 times for 15 min each in 0.02 M PBS. Sections were incubated overnight in 0.02 M PBS containing 0.3% Triton X100, 0.02% sodium azide and primary antibodies (**Table 1**) at room temperature in a humid chamber. In order to readily identify fine subcellular compartments expressing GFP/eGFP or GCaMP6f and to not miss any transgene expression, GFP/eGFP/GCaMP6f signal was amplified using an antibody directed against GFP. Of note, GFP/eGFP/GCaMP6f signal was visible without such amplification. The following day, sections were washed 3 times for 15 min each in 0.02 M PBS, and incubated for 2 h at room temperature with secondary antibodies diluted in 0.02 M PBS containing 0.3% Triton X100 and 0.02% sodium azide. Then, sections were washed 3 times for 15 min in 0.02 M PBS and mounted between slide and coverslip using Vectashield medium containing DAPI (Vector Laboratories). Negative controls, *i.e.* slices incubated with secondary antibodies only, were used to set criteria (gain, exposure time) for image acquisition in each experiment. Image acquisition was performed with an Axio Observer Z1 epifluorescence Zeiss microscope, an ORCA Flash 2.8 million pixel camera, and a PlanNeoFluar 20x/0.5NA objective. Images were extracted using the ZEN 2011 blue edition software (Zeiss). Cell counting measurements were performed using ImageJ software (National Institutes of Health, USA). Because it was difficult to discriminate individual SGCs within the several SGCs surrounding a single neuronal cell body, “rings” surrounding individual neuronal cell bodies were quantified.

### Calcium imaging

Acute intact DRG preparations were prepared from GFAP-Cre::GCaMP6f mice. Briefly, vertebras and dura mater were removed and lumbar L4 and L5 DRGs were exposed and immediately covered with ice cold (slushy) incubation ACSF solution (95 mM NaCl, 1.9 mM KCl, 1.2 mM KH_2_PO_4_, 0.5 mM CaCl_2_, 7 mM MgSO_4_, 26 mM NaHCO_3_, 15 mM glucose, 50 mM sucrose) and bubbled with 95% O_2_ and 5% CO_2_. DRGs were incubated for 30 min at 35°C in the incubation solution and then left to recover for 1 h 30 min at room temperature. A single DRG was placed in the recording chamber of a custom-built 2-photon laser-scanning microscope with a 20x water immersion objective (x20/0.95w XLMPlanFluor, Olympus). GCaMP6f was excited at 920 nm with a Ti:Sapphire laser (Mai Tai HP; Spectra-Physics). DRGs were continuously superfused at a rate of 4 ml/min with recording solution (127 mM NaCl, 1.9 mM KCl, 1.2 mM KH_2_PO_4_, 2.4 mM CaCl_2_, 1.3 mM MgSO_4_, 26 mM NaHCO_3_, 15 mM glucose) and bubbled with 95% O2 - 5% CO2. To evoke intracellular Ca^2+^ elevations in DRG GCaMP6f-expressing cells, a cocktail of agonists to endogenous G_q_ GPCRs containing 50 µM DHPG (Abcam), 10 µM histamine (Sigma Aldrich), 10 µM carbachol (Abcam), and 50 µM ATP (Sigma Aldrich) was bath applied for 30 sec.

## Results

### TRANSGENIC MOUSE LINES FOR INVESTIGATING THE DISTRIBUTION AND MORPHOLOGY OF SGCs AND MACROPHAGES

#### S100β-eGFP mice exhibit eGFP expression in both SGCs and sensory neurons

S100β protein is a commonly used astrocytic marker in the brain and spinal cord and has also been reported to be a marker of SGCs in DRGs (12, 32). The S100β-eGFP transgenic mouse (5), which expresses enhanced GFP (eGFP) under the control of the S100β promoter, has proved to be a useful tool to selectively label almost all astrocytes (32, 33). In the primary visual cortex (V1), we found that eGFP was expressed in 86% of astrocytes (**Fig 1A**; **Table 2**) and only marginally (0.02%) in neurons (**Fig 1B**; **Table 2**), corroborating those previous studies. This mouse line appears also to be valuable for labeling SGCs within DRGs, since eGFP immunoreactivity was detected in 85.8% of SGCs as shown by the colocalization of eGFP with GLAST, a specific SGC marker (**Fig 2A**; **Table 2**). In addition, some cells expressing eGFP, and exhibiting a typical feature of non-myelinating Schwann cells, were found in nerves attached to DRGs (**Fig 2C**; **Table 2**); this is expected for cells known to synthetize S100β (34, 35). However, 13.5% of sensory neuron soma (**Fig 2B**; **Table 2**) and their corresponding axons (**Fig 2C**) were found to express eGFP. To test whether this unexpected neuronal eGFP expression reflected normal endogenous S100β protein expression in DRGs of wildtype mice, we used an antibody directed against S100β. S100β endogenous protein was detected in both DRG SGCs and neurons of wildtype mice (**S1 Fig**), in agreement with the eGFP expression pattern observed in S100β-eGFP mice. Hence, the S100β promoter does not represent a valuable promoter to target transgene expression selectively in DRG SGCs. However, it does allow eGFP expression in the vast majority of DRG SGCs.

**Figure 1.**
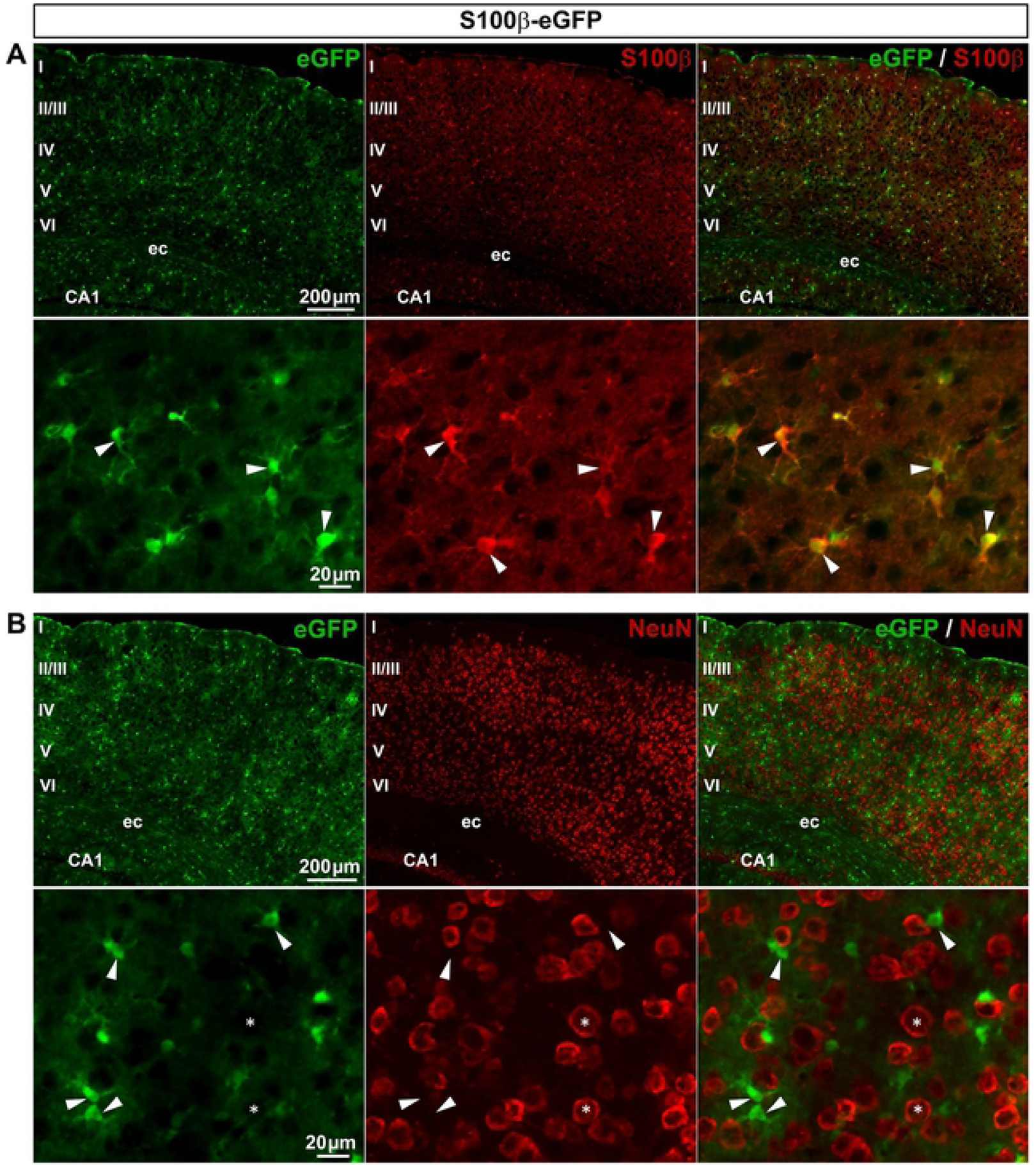
Expression of eGFP in S100β-eGFP mouse brain. **A**, **B**, Representative images of immunohistochemistry in the primary visual cortex (V1) from S100β-eGFP mice showing eGFP-expressing cells (**A** & **B left**, green), S100β−expressing astrocytes (**A middle**, red, arrowheads), and NeuN-expressing neurons (**B middle**, red, asterisks). **A** & **B right** show superimposed pictures. In **A** & **B**, bottom panel images correspond to enlargements of top panel images. For each row, scale bar in left picture applies to middle and right corresponding pictures. Abbreviations: ec, external capsule; CA1, subfield 1 of Ammon’s horn; I-VI, layers of V1.

**Figure 2.**
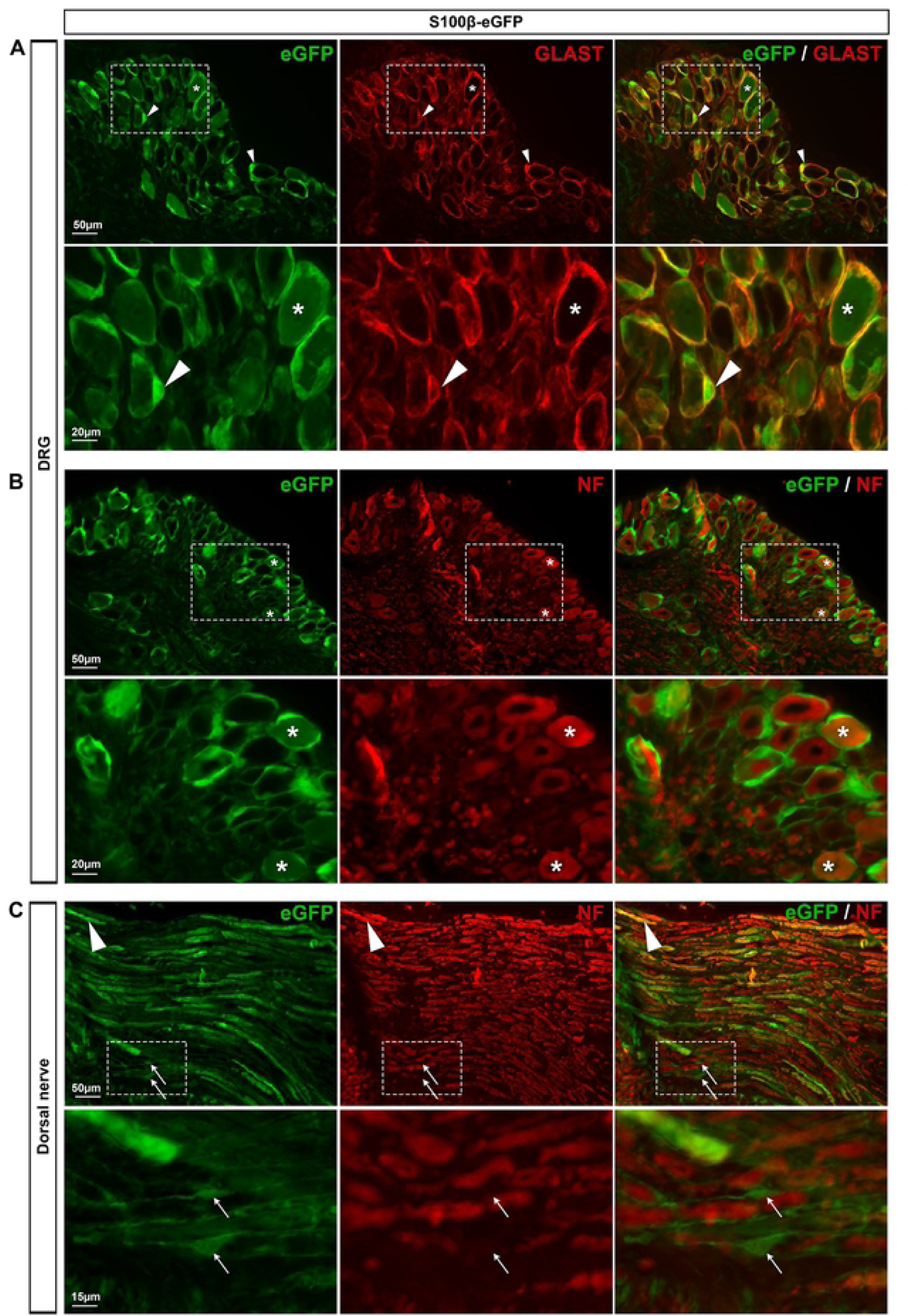
Expression of eGFP in S100β-eGFP mouse DRGs. **A**, **B**, Representative images of immunohistochemistry in DRGs from S100β-eGFP mice showing eGFP-expressing cells (**A-C left**, green), GLAST-expressing SGCs (**A middle**, red, arrowheads), and neurofilament-expressing neurons (**B middle**, red, asterisks). **C**, Images of proximal DRG nerves showing eGFP staining (**C left**, green), non-myelinating Schwann cells (**C left**, arrows) and neurofilament-expressing axons (**C middle**, arrowheads). **A-C right** show superimposed pictures. In **A**-**C**, bottom panel images correspond to enlargements of boxed areas in top panel images. For each row, scale bar in left picture applies to middle and right corresponding pictures.

#### ALDH1L1-eGFP mice express eGFP in a subset of SGCs and some neurons

Another recently discovered marker of astrocytes is the protein aldehyde dehydrogenase 1 family member L1 [aldh1l1; (2)]. The ALDH1L1 promoter has been used to generate several transgenic mouse lines (2,8,36), including the ALDH1L1-eGFP mice that exhibit eGFP selectively in most astrocytes of the CNS (2). In agreement, we found that eGFP is expressed in 82.1% of V1 astrocytes (**Fig 3A**; **Table 2**) with no detectable expression in neurons (**Fig 3B**; **Table 2**). The distribution of eGFP-expressing cells was then investigated in DRGs of ALDH1L1-eGFP mice. Enhanced GFP immunoreactivity was observed only in 56.4% of SGCs (**Fig 4A**; **Table 2**) as well as in a low percentage (7.2%) of sensory neuron soma, making this transgenic line unsuitable for the specific visualization of the majority of SGCs.

**Figure 3.**
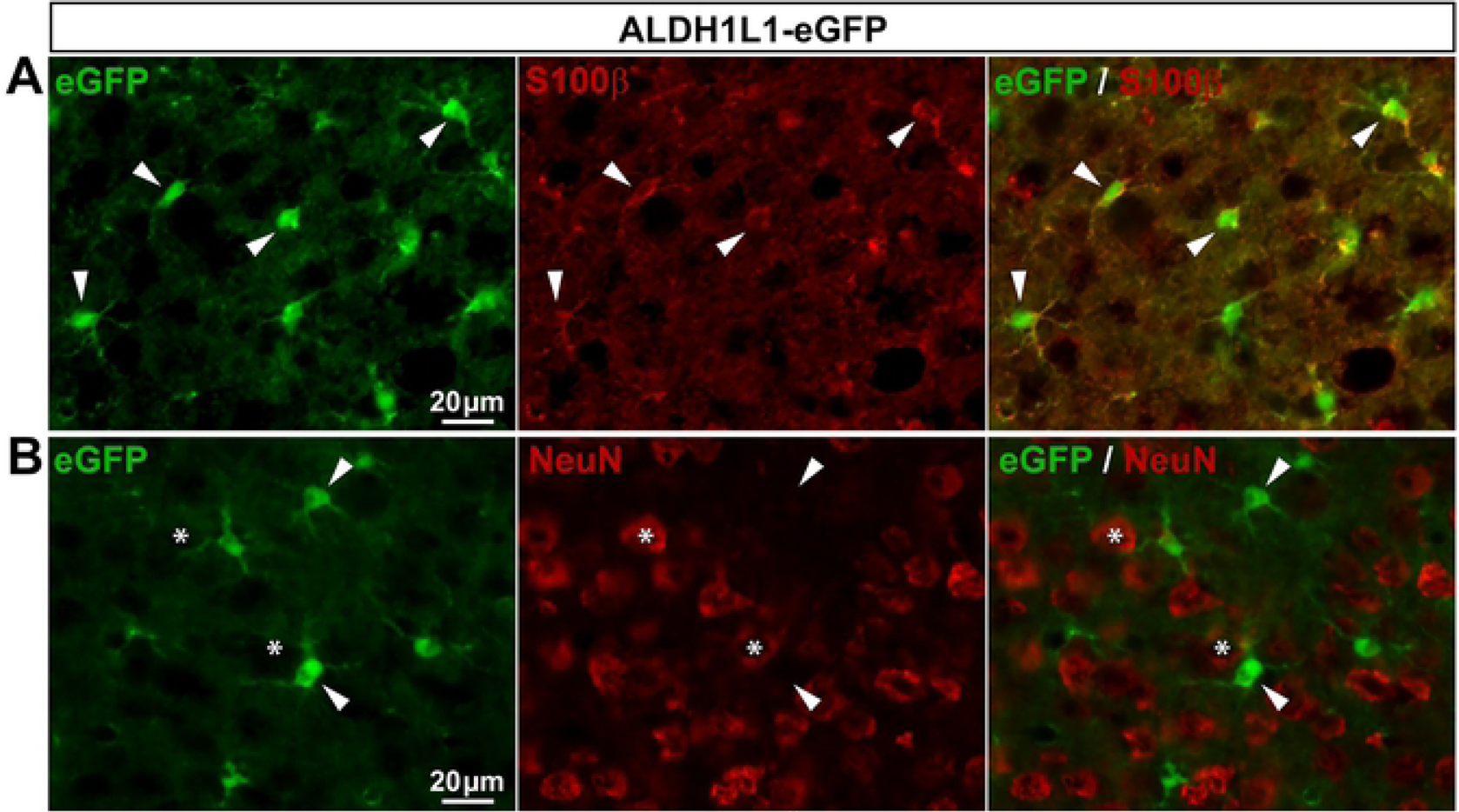
Expression of eGFP in ALDH1L1-eGFP mouse brain. **A**, **B**, Representative images of immunohistochemistry in V1 from ALDH1L1-eGFP mice showing eGFP-expressing cells (**A** & **B left**, green), S100β−expressing astrocytes (**A middle**, red, arrowheads), and NeuN-expressing neurons (**B middle**, red, asterisks). **A** & **B right** show superimposed pictures. For each row, scale bar in left picture applies to middle and right corresponding pictures.

**Figure 4.**
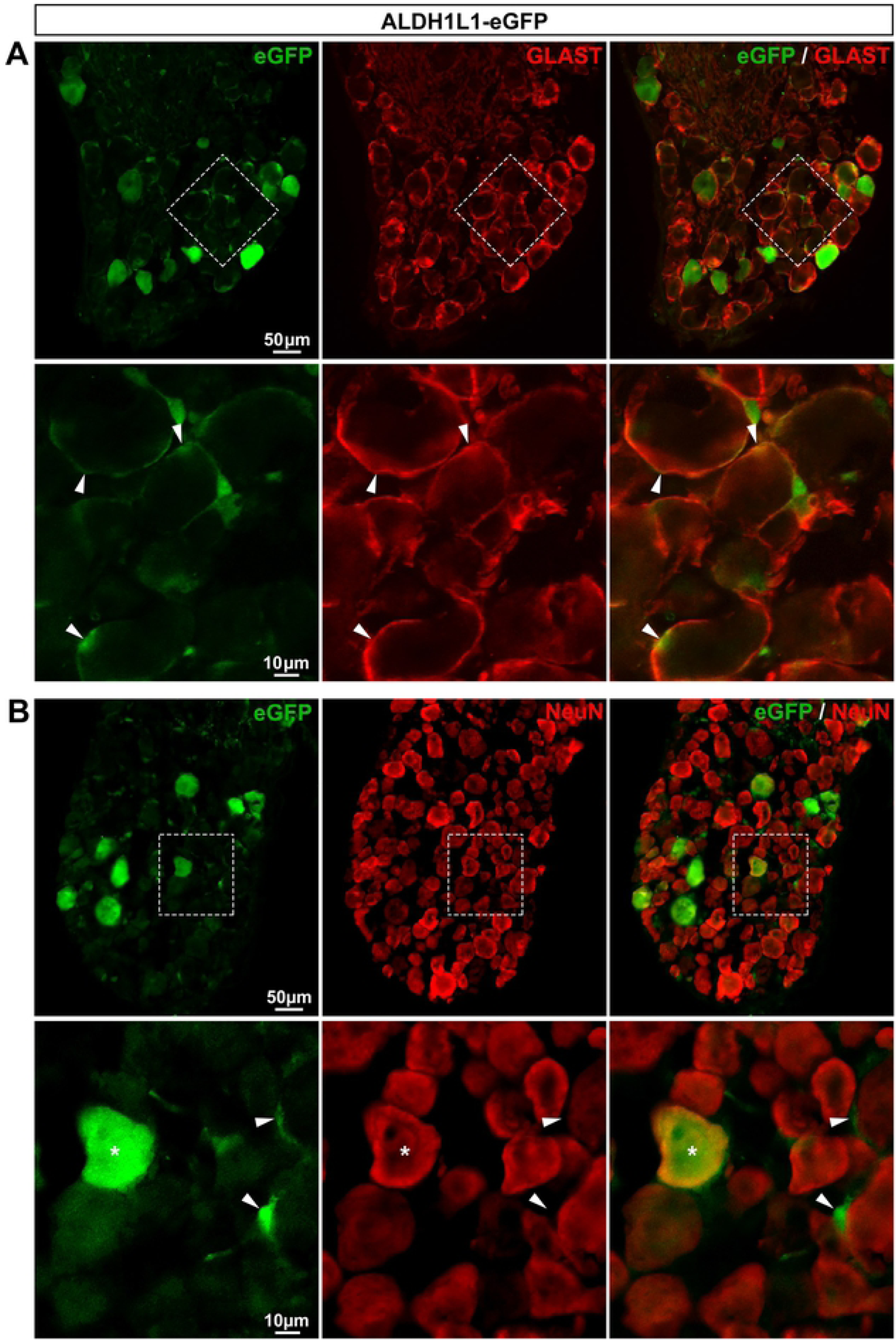
Expression of eGFP in ALDH1L1-eGFP mouse DRGs. **A**, **B**, Representative images of immunohistochemistry in DRGs from ALDH1L1-eGFP mice showing eGFP-expressing cells (**A** & **B left**, green), GLAST-expressing SGCs (**A middle**, red, arrowheads), and NeuN-expressing neurons (**B middle**, red, asterisks). **A** & **B right** show superimposed pictures. In **A** & **B**, bottom panel images correspond to enlargements of boxed areas in top panel pictures. In A bottom row, boxed area has been rotated by about 45° clockwise. For each row, scale bar in left picture applies to middle and right corresponding pictures.

#### CX3CR1-eGFP mice show specific eGFP expression in macrophages

The CX3C chemokine receptor 1 (CX3CR1), known as the fractalkine receptor (3), is a marker of microglial cells (37, 38). The CX3CR1-eGFP mouse line, expressing eGFP under the control of the CX3CR1 promoter (3), has been extremely useful to visualize microglial cells and dynamic changes in their morphology. In support of this previous study, we found a selective eGFP expression in 99.5% of V1 microglial cells with no detectable expression in neurons (**Fig 5A,B**; **Table 3**). Since microglial cells are the CNS resident macrophages, we reasoned that peripheral macrophages should also express eGFP in DRGs from CX3CR1-GFP mice. Indeed, we observed that eGFP was expressed in 90.9% of Iba1-expressing macrophages (**Fig 6A**; **Table 3**), validating the use of the CX3CR1-eGFP mouse line for labeling a prominent number of DRG macrophages and investigating their morphological remodeling. Note that a few eGFP-positive elements did not express Iba1 (**Fig 6A**, arrow). Because Iba1-immunopositive profiles are spotty and not found within the whole cytosol, but instead are localized in some subcellular compartments of macrophages, such eGFP-positive elements might correspond to Iba1-negative macrophage compartments.

**Figure 5.**
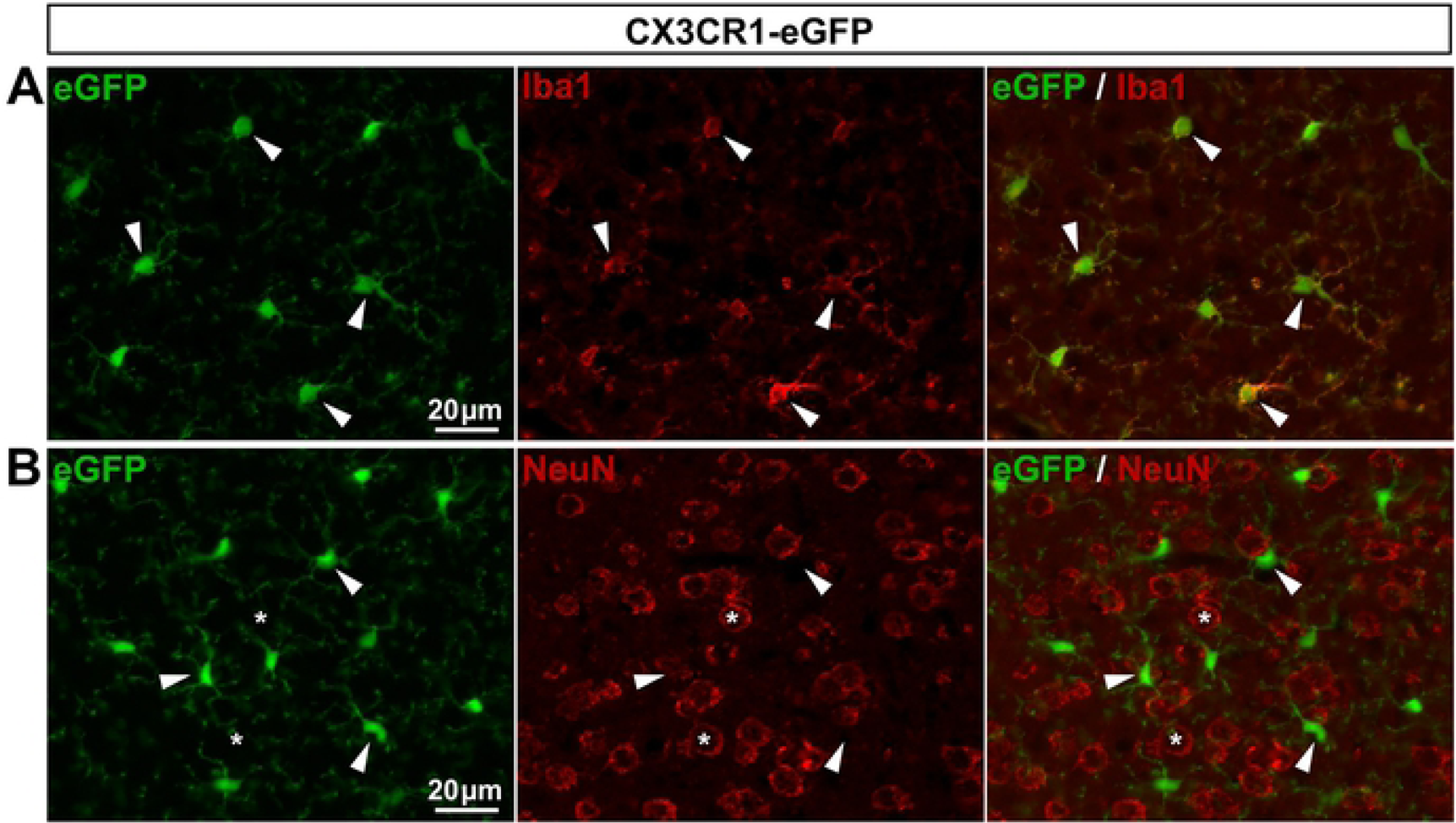
Expression of eGFP in CX3CR1-eGFP mouse brain. **A**, **B**, Representative images of immunohistochemistry in V1 from CX3CR1-eGFP mice showing eGFP-expressing cells (**A** & **B left**, green), Iba1-expressing microglial cells (**A middle**, red, arrowheads), and NeuN-expressing neurons (**B middle**, red, asterisks). **A** & **B right** show superimposed pictures. For each row, scale bar in left picture applies to middle and right corresponding pictures.

**Figure 6.**
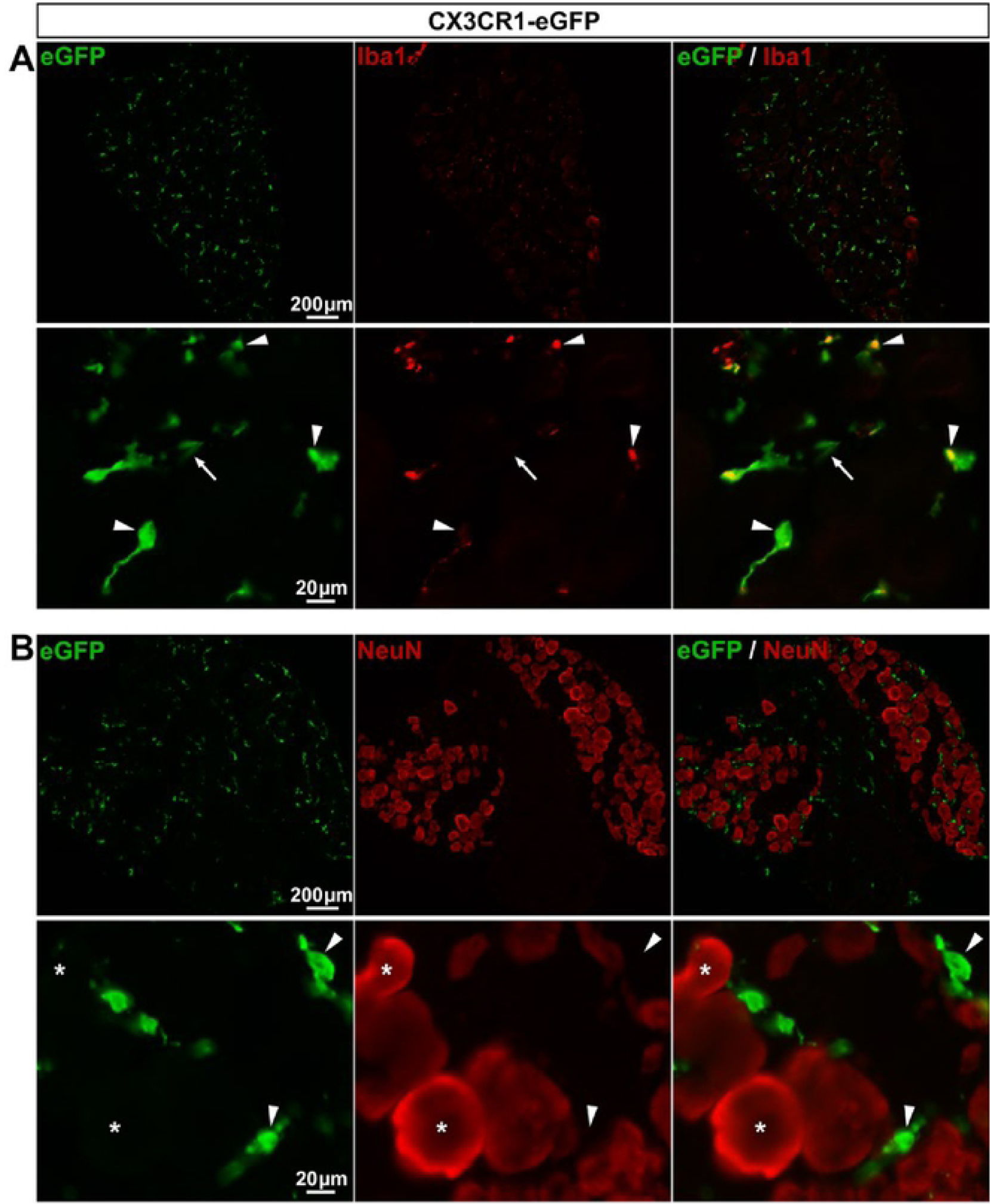
Expression of eGFP in CX3CR1-eGFP mouse DRGs. **A**, **B**, Representative images of immunohistochemistry in DRGs from CX3CR1-eGFP mice showing eGFP staining (**A** & **B left**, green, arrowheads), Iba1 staining (**A middle**, red, arrowheads), and neurons identified by NeuN immunoreactivity (**B middle**, red, asterisks). In **A bottom panel**, the arrow points to an eGFP-expressing element that does not colocalize with Iba1 staining. **A** & **B right** show superimposed pictures. In **A** & **B**, bottom panel images correspond to enlargements of top panel pictures. For each row, scale bar in left picture is applied to middle and right corresponding pictures.

### MOUSE LINES FOR INVESTIGATING THE ROLES OF SGC Ca^2+^ SIGNALING

#### GFAP-Cre::GCaMP6f mouse line expresses GCaMP6f Ca^2+^ biosensor mainly in DRG sensory neurons

Emerging evidence showing that Ca^2+^ is an important signaling messenger in SGCs (12,39,40) prompted us to search for molecular tools to probe SGC Ca^2+^ dynamics. To do so, we took advantage of the powerful CAG-lox-STOP-lox-GCaMP6f transgenic mouse line (7) that was recently developed to express the genetically-encoded Ca^2+^ indicator GCaMP6f in cell types of interest. We first crossed CAG-lox-STOP-lox-GCaMP6f mice with GFAP-Cre mice (29) to generate GFAP-Cre::GCaMP6f double transgenic mice. GFAP is indeed extensively used as a gold standard astrocytic marker, which expression is enhanced in reactive astrocytes during aging, CNS injury, pain, and diseases (41). GFAP has also been reported as a marker of DRG SGCs (11), although we have not found convincing immunohistochemical evidence in the literature showing clear SGC GFAP expression under physiological conditions. In agreement, in our hands, immunohistochemical experiments conducted on DRGs from wildtype mice using two different antibodies directed against GFAP (**Table 1**) have revealed almost no GFAP-expressing SGCs (data not shown). However, a large number of studies have shown increases in SGC GFAP expression level under pathological conditions, including, peripheral nerve injury, DRG compression and pain (17, 24).

In the CNS, GFAP is known to be expressed in neural progenitors during developmental stages, giving rise to neurons, astrocytes and oligodendrocytes, thus preventing the use of the non-inducible GFAP-Cre mice and Cre-LoxP system to selectively study astrocytes (29,42–44). The observation that astrocytes, neurons, and neuropil from V1 GFAP-Cre::GCaMP6f expressed GCaMP6f (**Fig 7A,B**) was consistent with those previous studies and the view that recombination occurs during development. However, because SGCs derive from the neural crest, a different lineage than astrocytic lineage, there was a possibility that GFAP-Cre mice could be useful to drive gene expression selectively in a great proportion of SGCs. In DRGs from GFAP-Cre::GCaMP6f mice, GCaMP6f was expressed in only 1.8% of SGCs while, to our surprise, it was observed in 58.5% of sensory neurons (including small- and large-sized diameter neurons; **Fig 8A**; **Table 2**). To assess whether GCaMP6f was expressed under detectable levels in SGCs, SGC functional Ca^2+^ signals were registered using 2-photon imaging in intact *ex vivo* DRGs. Bath application of an agonist cocktail to G_q_ GPCRs did not evoke any Ca^2+^ elevations in cells morphologically identified as SGCs (*i.e*. ring-shaped cells surrounding neuronal soma). However, 23.5% of GCaMP6f-expressing sensory neurons exhibited marked intracellular Ca^2+^ elevations (**Fig 8B**). Taken together these results show that the GFAP-Cre::GCaMP6f double transgenic mouse line is not an adequate tool for studying Ca^2+^ dynamics selectively in DRG SGCs.

**Figure 7.**
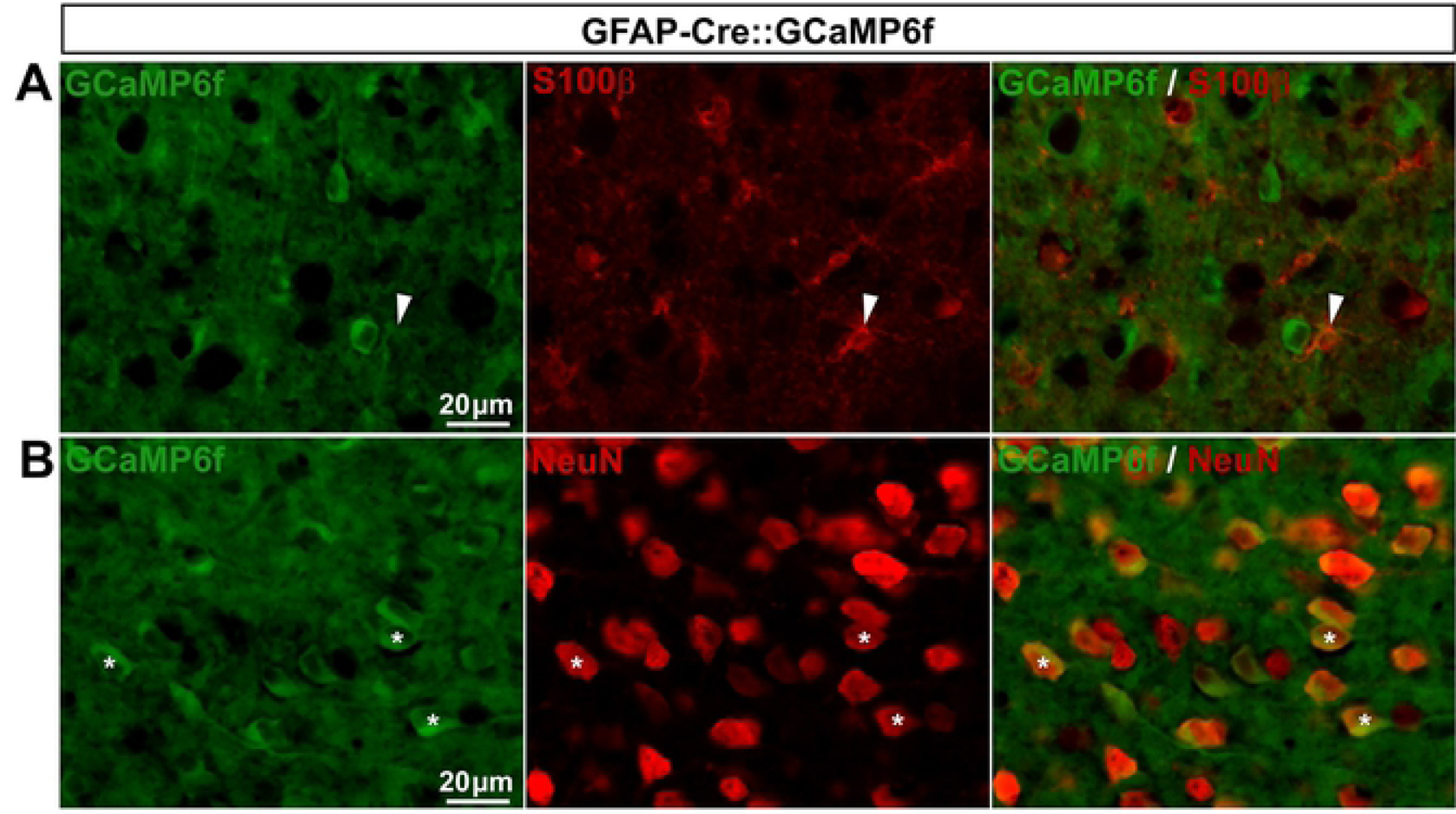
Expression of GCaMP6f in GFAP-Cre::GCaMP6f mouse brain. **A**, **B**, Representative images of immunohistochemistry in V1 from GFAP-Cre::GCaMP6f mice showing GCaMP6f ubiquitous expression (**A** & **B left**, green), S100β−expressing astrocytes (**A middle**, red, arrowheads), and NeuN-expressing neurons (**B middle**, red, asterisks). **A** & **B right** show superimposed pictures. For each row, scale bar in left picture applies to middle and right corresponding pictures.

**Figure 8.**
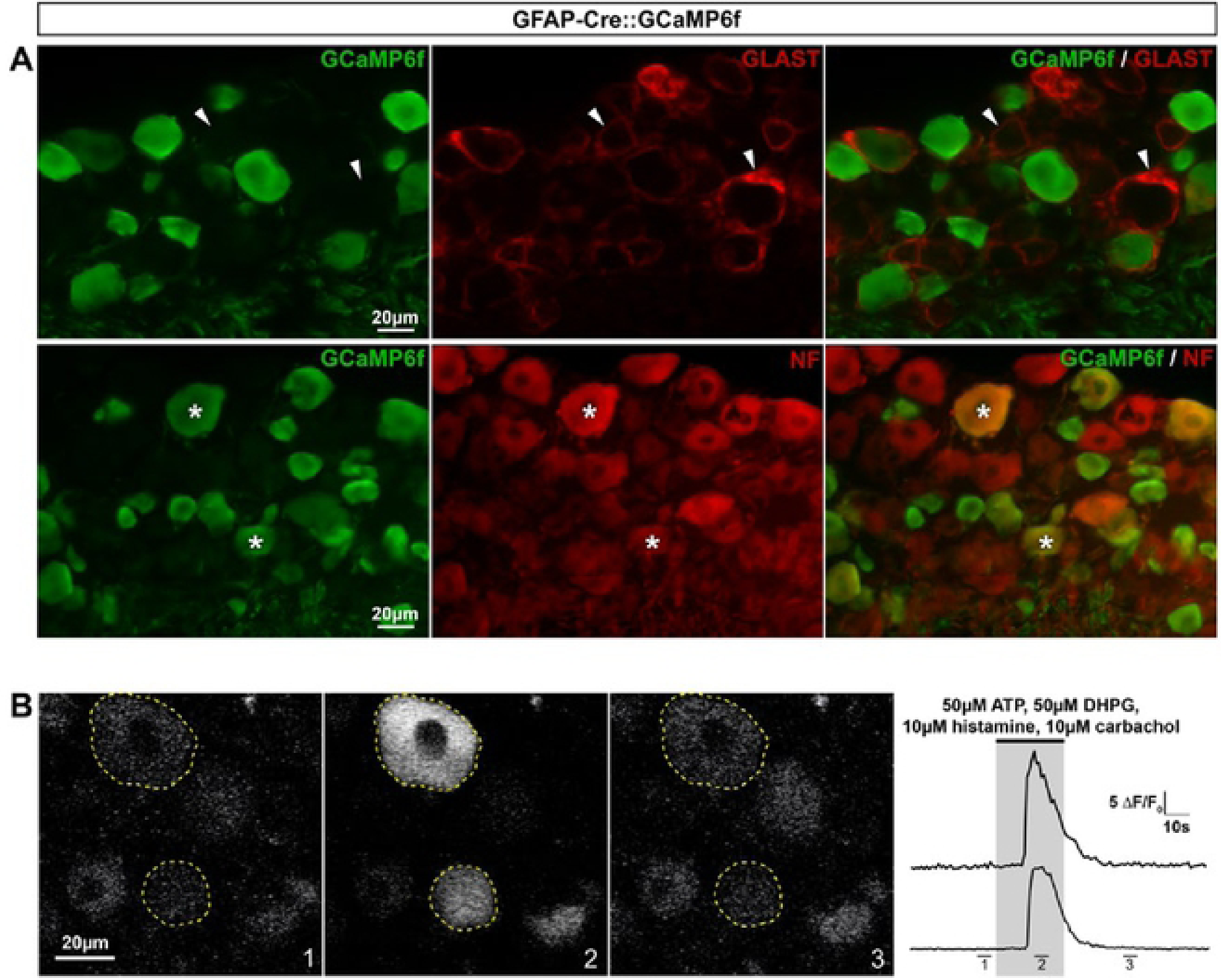
Cellular expression and functionality of GCaMP6f in GFAP-Cre::GCaMP6f mouse DRGs. **A**, Representative images of immunohistochemistry in DRGs from GFAP-Cre::GCaMP6f mice showing GCaMP6f staining (**top** & **bottom left panels**, green), GLAST-expressing SGCs (**top middle panel**, red, arrowheads), and small and large sensory neurons (**bottom middle panel**, red, asterisks). **Top** & **bottom right panels** show superimposed pictures. For each row, scale bar in left picture applies to middle and right corresponding pictures. **B**, Representative images of 2-photon Ca^2+^ imaging experiment in *ex vivo* DRGs where neuronal GCaMP6f-expressing cell bodies (outlined areas of interest, **left panel**) exhibit intracellular Ca^2+^ increases; ① baseline, ② G_q_ GPCR agonist cocktail (50µM ATP, 10µM Histamine, 10µM Carbachol and 50µM DHPG) application, and ③ wash (**right panel**).

#### GLAST-CreERT2::GCaMP6f and Cx30-CreERT2::GCaMP6f mice express GCaMP6f in a few or no SGCs, respectively

GLAST has been conventionally used to identify a subset of astrocytes and the majority of DRG SGCs (45–47), making the GLAST promoter a good candidate to target gene expression in a large number of SGCs. Additionally, connexin 30 (Cx30) has also been used as a marker of a fraction of astrocytes (45), although its expression in DRG SGCs has not yet been reported. To determine whether GLAST and Cx30 promoters could drive GCaMP6f expression selectively in SGCs, we crossed both inducible GLAST-CreERT2 (28) and Cx30-CreERT2 (28) mice with CAG-lox-STOP-lox-GCaMP6f mice. To induce GCaMP6f expression in the resultant GLAST-CreERT2::GCaMP6f and Cx30-CreERT2::GCaMP6f double transgenic lines, mice were treated with 1 mg tamoxifen per day during 5 consecutive days. As expected, in the CNS (V1), GCaMP6f was detected in 56.3% and 46.4% of astrocytes, in these two lines respectively (**Fig 9A**; **Fig 10A**; **Table 2**) with an insignificant expression in neurons (0.03% and 0.1%, respectively; **Fig 9B**; **Fig 10B**; **Table 2**). In contrast, DRGs from GLAST-CreERT2::GCaMP6f mice showed GCaMP6f immunoreactivity in only 5.4% of SGCs and a low percentage (3.8%) of sensory neurons (**Fig 11A,B**; **Table 2**), invalidating this mouse line for investigating Ca^2+^ dynamics selectively in SGCs. Furthermore, no GCaMP6f expression at all was found in SGCs or sensory neurons of Cx30-CreERT2::GCaMP6f (**Fig 12A,B**; **Table 2**), making this mouse line unsuitable to study Ca^2+^ signaling in DRG SGCs. Surprisingly though, GCaMP6f was observed in non-identified cells, which occasionally expressed macrophage markers (**Fig 12C**), suggesting that a few of them are macrophages. Of note, GCaMP6f was also found to be expressed at the DRG surface, possibly in tissues encapsulating DRGs (**Fig 12B**).

**Figure 9.**
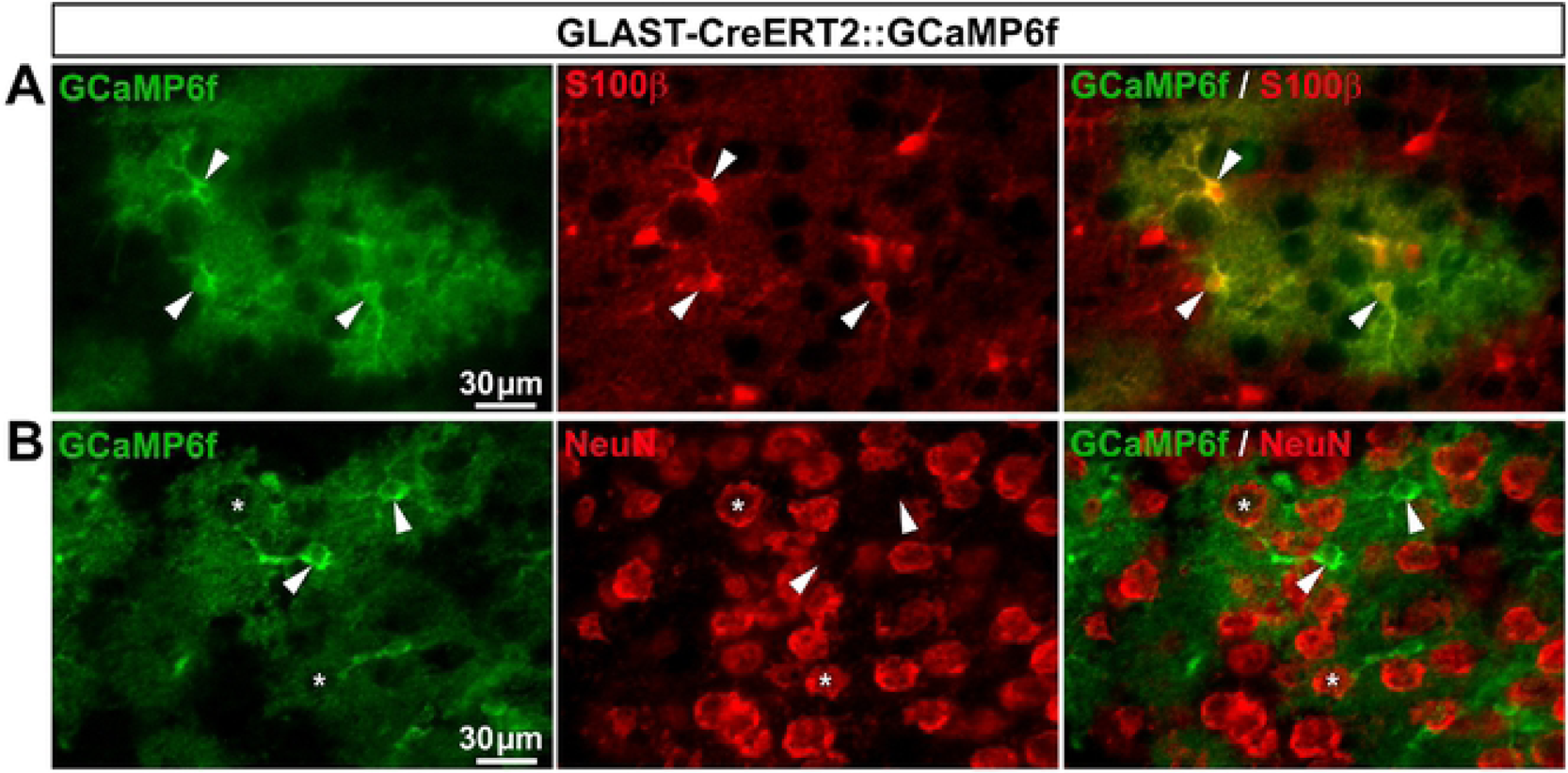
Expression of GCaMP6f in GLAST-CreERT2::GCaMP6f mouse brain. **A**, **B**, Representative images of immunohistochemistry in V1 from GLAST-CreERT2::GCaMP6f mice showing GCaMP6f-expressing cells (**A** & **B left**, green, arrowheads), S100β−expressing astrocytes (**A middle,** red, arrowheads), and NeuN-expressing neurons (**B middle**, red, asterisks). **A** & **B right** show superimposed pictures. For each row, scale bar in left picture applies to middle and right corresponding pictures.

**Figure 10.**
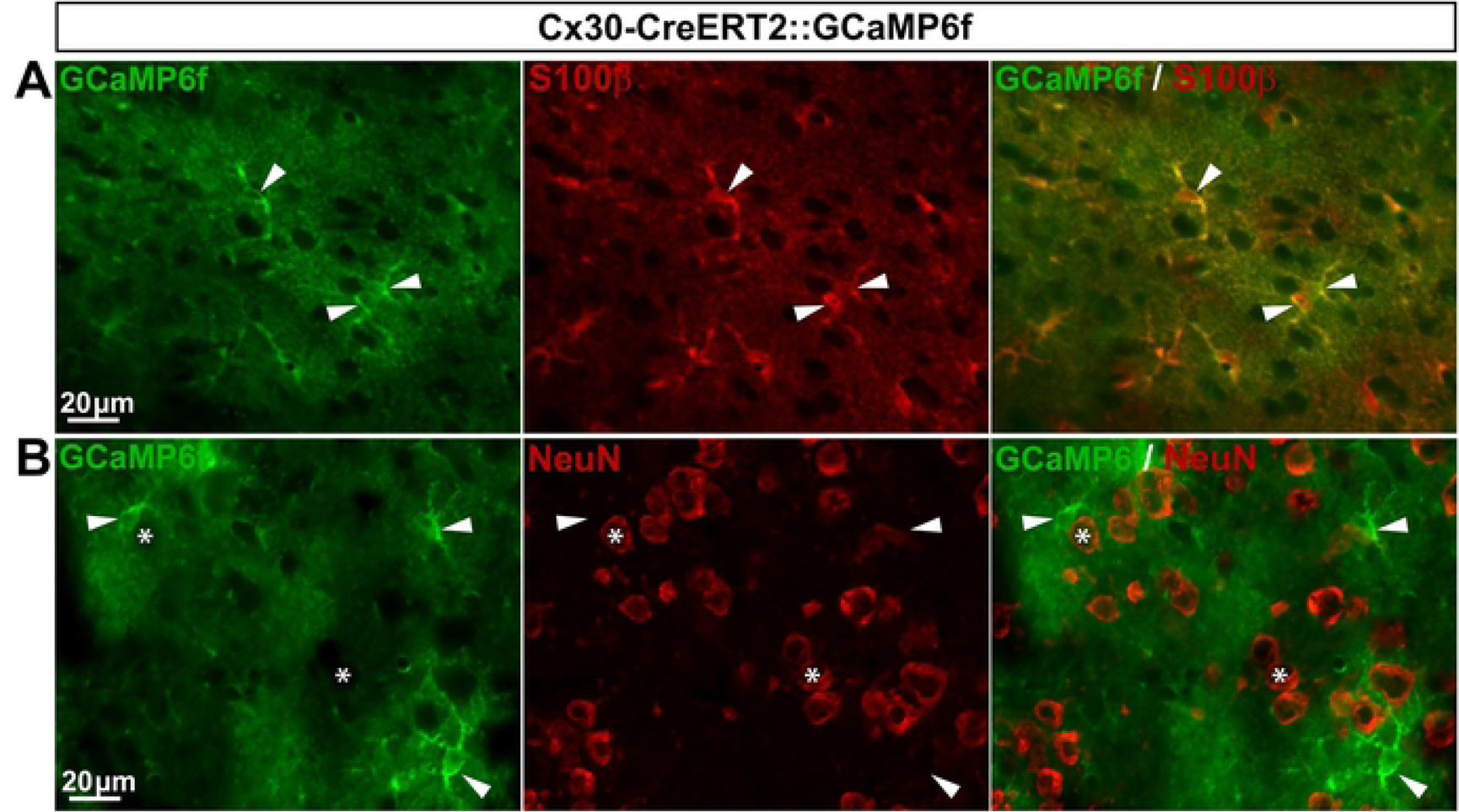
Expression of GCaMP6f in Cx30-CreERT2::GCaMP6f mouse brain. **A**, **B**, Representative images of immunohistochemistry in V1 from Cx30-CreERT2::GCaMP6f mice showing GCaMP6f-expressing cells (**A** & **B left**, green, arrowheads), S100β−expressing astrocytes (**A middle,** red, arrowheads), and NeuN-expressing neurons (**B middle**, red, asterisks). **A** & **B right** show superimposed pictures. For each row, scale bar in left picture applies to middle and right corresponding pictures.

**Figure 11.**
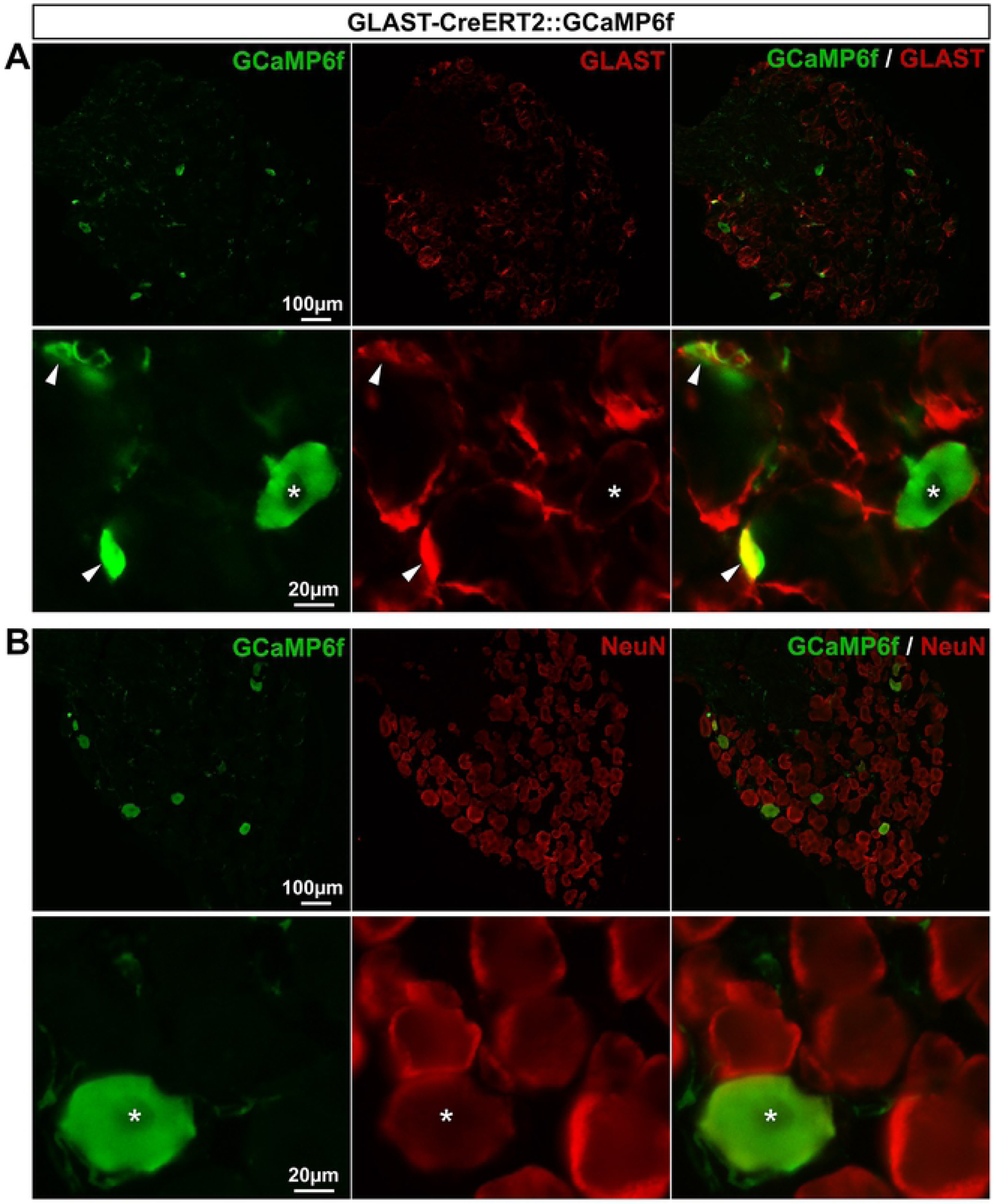
Expression of GCaMP6f in GLAST-CreERT2::GCaMP6f mouse DRGs. **A**, **B**, Representative images of immunohistochemistry in DRGs from GLAST-CreERT2::GCaMP6f mice showing GCaMP6f labeling (**A** & **B left**, green), GLAST-expressing SGCs (**A bottom panel**, red, arrowheads,), and NeuN-expressing neurons (**B bottom panel**, red, asterisks). **A** & **B right** show superimposed pictures. In **A** & **B**, bottom panel images correspond to enlargements of top panel pictures. For each row, scale bar in left picture applies to middle and right corresponding pictures.

**Figure 12.**
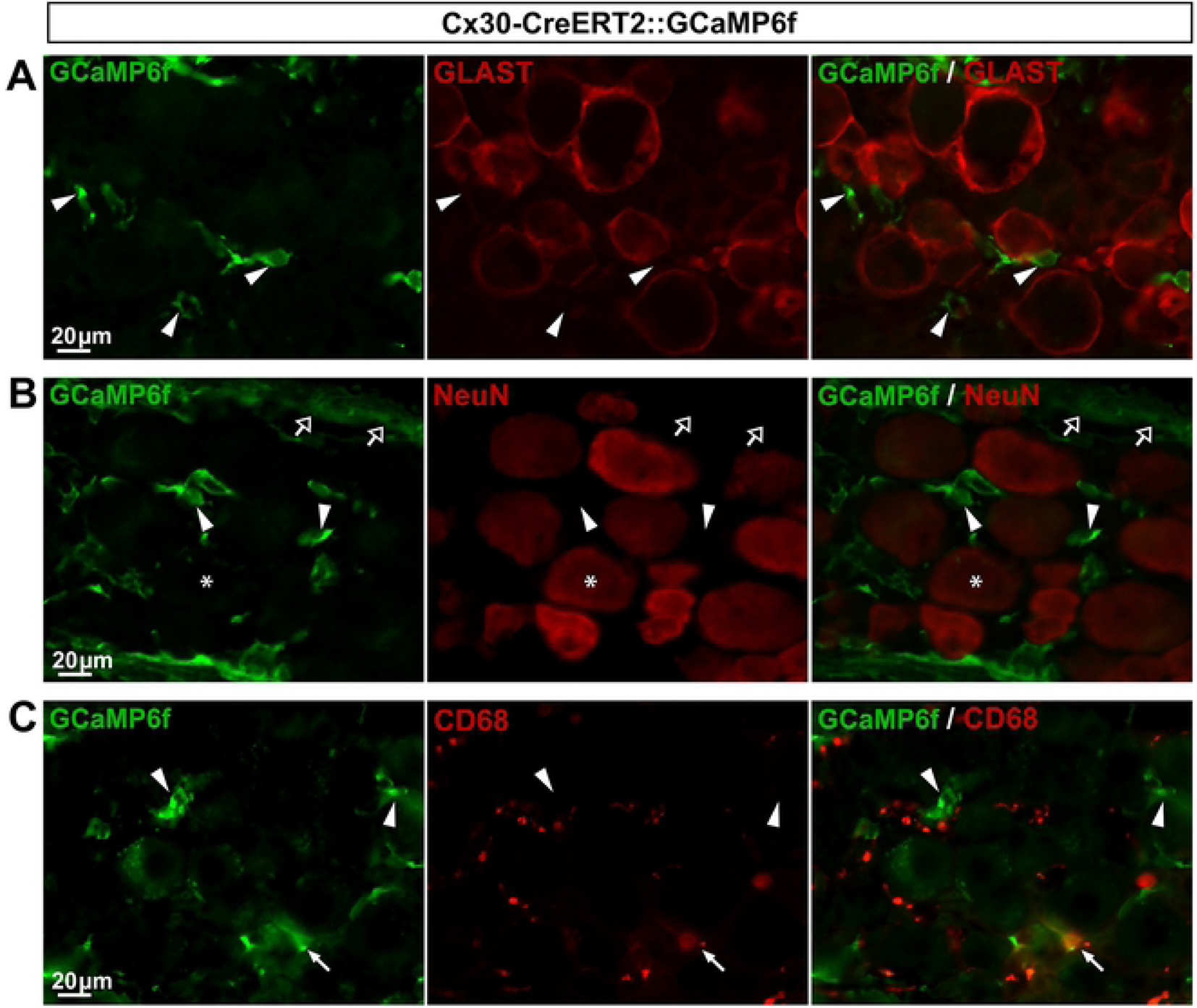
Expression of GCaMP6f in Cx30-CreERT2::GCaMP6f mouse DRGs. **A**-**C**, Representative images of immunohistochemistry in DRGs from Cx30-CreERT2::GCaMP6f mice showing GCaMP6f immunoreactivity (**A–C left**, green, arrowheads), GLAST-expressing SGCs (**A middle**, red), and NeuN-expressing neurons (**B middle**, red, asterisks). In **B left**, empty arrows point to GCaMP6f expression at the surface of the DRG. In **C left**, white arrow points to GCaMP6f staining that colocalizes with the macrophage marker CD68 immunoreactivity (**C middle**, red,). **A-B right** show superimposed pictures. For each row, scale bar in left picture applies to middle and right corresponding pictures.

#### Cx43-CreERT2::GCaMP6f mouse line allows the detection of Ca^2+^ transients in the vast majority of SGCs

Cx43 is a widely used specific marker of a large proportion of CNS astrocytes and almost all DRG SGCs (13, 48), making it particularly relevant to astrocyte and SGC research. In another attempt to establish a tool for monitoring Ca^2+^ transients selectively in a great number of SGCs, we generated the inducible Cx43-CreERT2::GCaMP6f double transgenic mouse line by crossing the tamoxifen-inducible Cx43-CreERT2 (30) with the CAG-lox-STOP-lox-GCaMP6f mice (7). Two weeks after tamoxifen treatment, V1 cortex and DRGs were analyzed. To our disappointment, GCaMP6f immunoreactivity was observed in only 9.5% of V1 astrocytes (**Fig 13A**; **Table 2**), a much lower percentage than previously reported in hippocampal astrocytes (∼70%; 31). Possible explanations for these discrepancies are regional (V1 *versus* hippocampus) variability in transgene recombination efficiency, in addition to the fact that we tested our mice 2 weeks after tamoxifen treatment compared to 4 weeks (31). Furthermore, GCaMP6f was also detectable in a small subset (4.4%) of neurons (**Fig 13B**; **Table 2**). These results suggest that Cx43-CreERT2 mouse line may be ineffective for driving substantial GCamP6f expression specifically in V1 astrocytes, even when treating mice 10 times (10 x 1 mg/kg tamoxifen) as previously reported (31).

**Figure 13.**
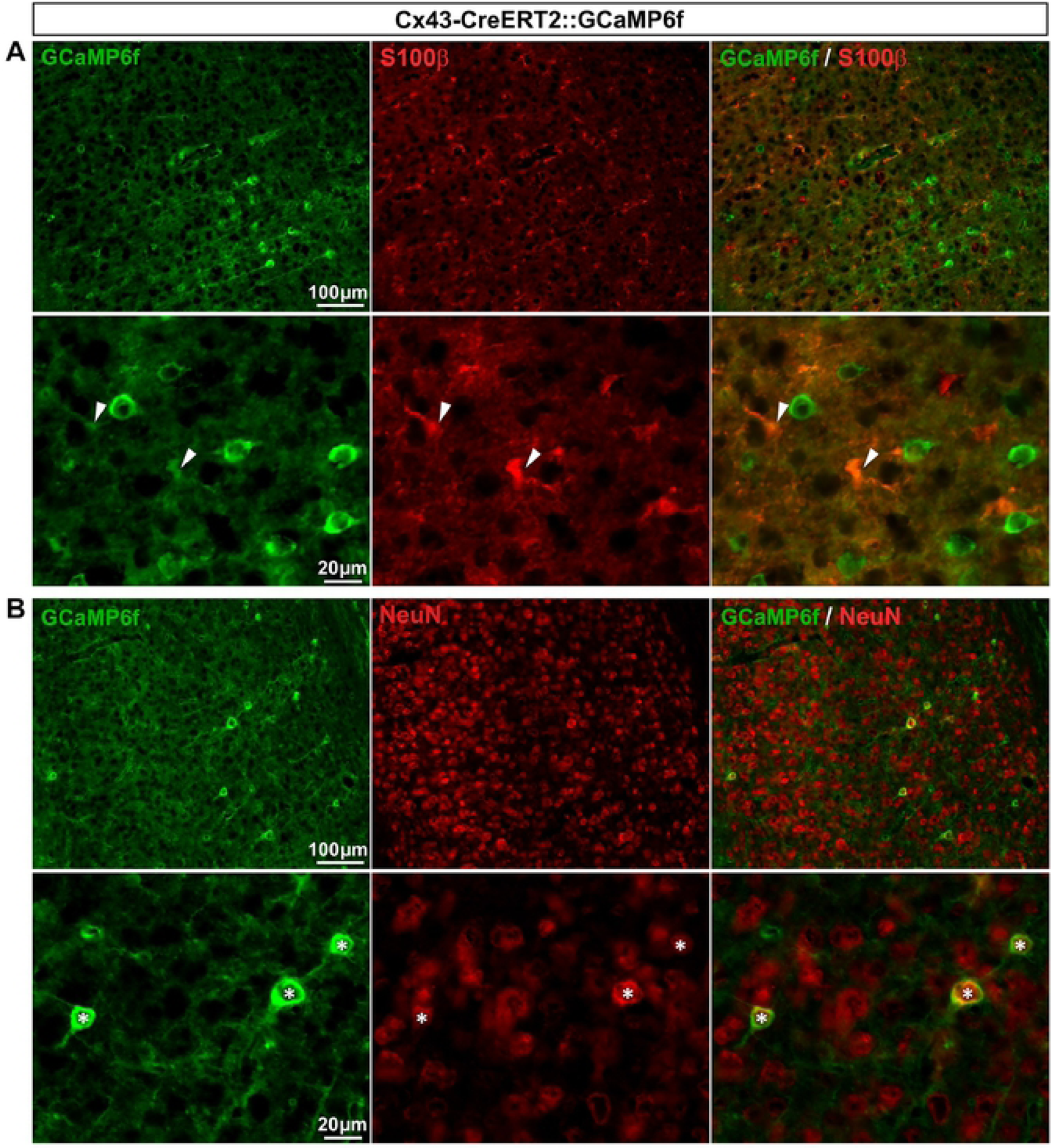
Expression of GCaMP6f in Cx43-CreERT2::GCaMP6f mouse brain. **A**, **B**, Representative images of immunohistochemistry in V1 from Cx43-CreERT2::GCaMP6f mice showing GCaMP6f immunoreactivity (**A** & **B left**, green), S100β−expressing astrocytes (**A middle bottom panel**, red, arrowheads) and NeuN-expressing neurons (**B bottom panel**, red, asterisk). **A** & **B right** show superimposed pictures. In **A** & **B**, bottom panel pictures correspond to enlargements of top panel pictures. For each row, scale bar in left picture applies to middle and right corresponding pictures.

However, and importantly, cellular immunohistochemical characterization in DRGs revealed that GCaMP6f was expressed in 92.6% of SGCs and a very small subset (4%) of neurons (**Fig 14A,B**; **Table 2**). In conclusion, the inducible Cx43-CreERT2::GCaMP6f line can be used to study the role of Ca^2+^ activity in the majority of DRG SGCs. We believe that this mouse line will prove to be a valuable tool to examine DRG SGC functions.

**Figure 14.**
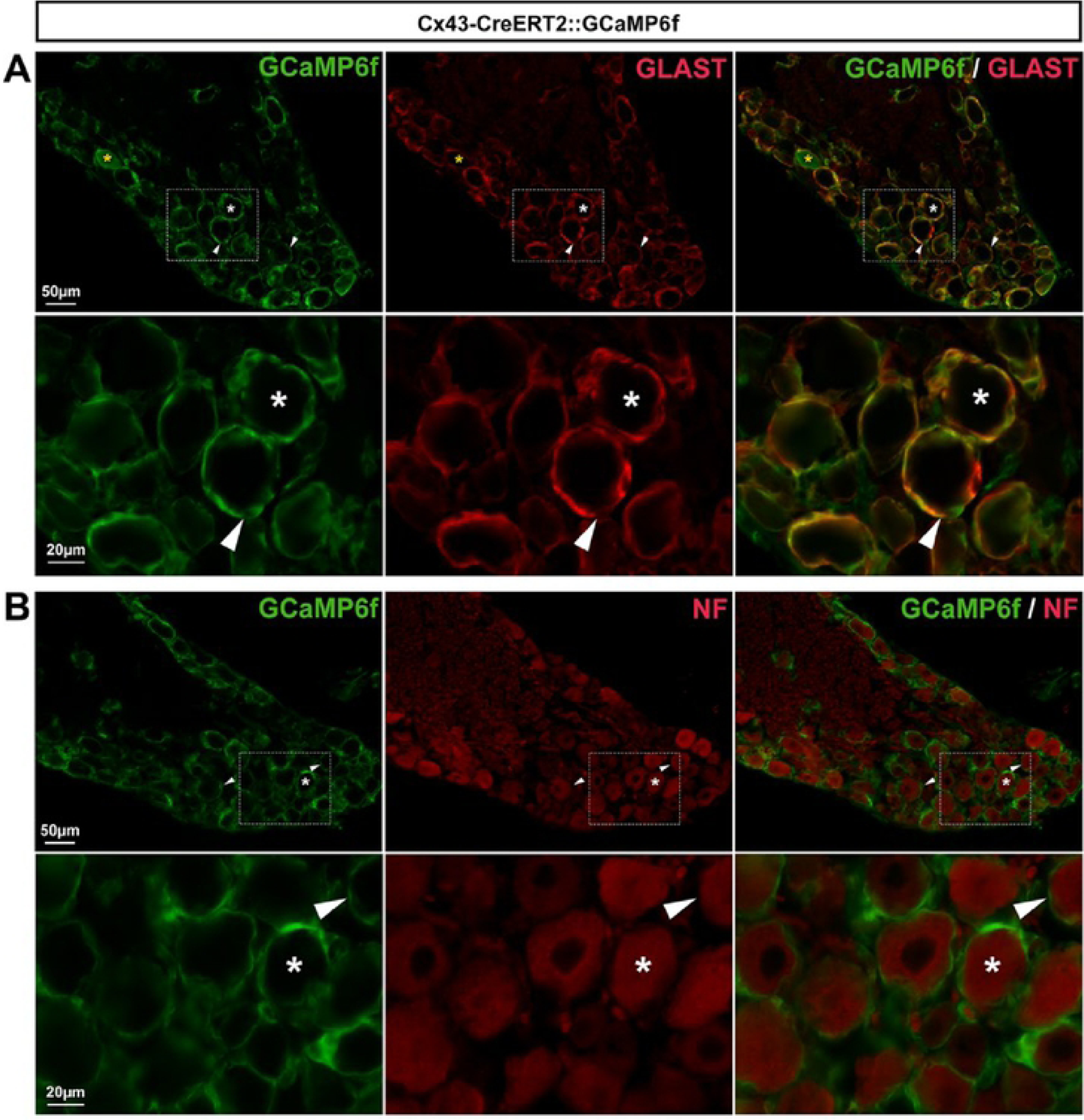
Expression of GCaMP6f in Cx43-CreERT2::GCaMP6f mouse DRGs. **A**, **B**, Representative images of immunohistochemistry in DRGs from Cx43-CreERT2::GCaMP6f mice showing GCaMP6f immunoreactivity (**A** & **B left**, green), GLAST-expressing SGCs (**A middle**, red, arrowheads), and NF-expressing neuronal cell bodies (**B middle**, red, asterisks). Note: Yellow asterisk (**A**, **top panel**) shows a single neuron expressing GCaMP6f. **A** & **B right** show superimposed pictures. In **A** & **B**, bottom panel pictures correspond to enlargements of boxed areas in top panel images. For each row, scale bar in left picture applies to middle and right corresponding pictures.

## Discussion

In this study we report the characterization of several tools for investigating SGC and macrophage morphological changes as well as Ca^2+^ activity within DRGs. Our data show that most tested transgenic mice widely used to investigate astrocyte morphology and function are not suitable for studying DRG SGCs. Indeed, among these mice, some exhibit ectopic transgene expression in small to large proportions of neurons while others show low to no transgene expression in SGCs.

However, we generated and identified a double transgenic line, named Cx43-CreERT2::GCaMP6f, allowing inducible GCaMP6f expression primarily in the vast majority of DRG SGCs (92.6%) with only a very small percentage (4%) of neurons expressing GCaMP6f. Considering the high GCaMP6f expression level detected in SGCs, ectopic CreERT2-mediated recombination in neurons might be easily reduced by simply decreasing the number of tamoxifen injections (< 10 injections) as well as time post-treatment (< 15 days). With the emerging interest in SGC Ca^2+^ signaling in modulating nociceptive neuron activity (39,40,49), this new line should be applicable for investigating a wide array of questions in pain research. Furthermore, we identified a second mouse line, called CX3CR1-eGFP, displaying eGFP expression selectively in most DRG macrophages. This line is likely to be useful for the study of spinal cord injury in which abnormal pain strongly correlates with an increased number of DRG macrophages (50).

Among the other mouse lines we characterized, S100β-eGFP line may be considered applicable to some extent, even though eGFP was expressed in a substantial percentage of sensory neurons (13.5%), but also in a great proportion of DRG SGCs (85.8%). The readily identifiable ring-shaped SGCs can indeed be almost undoubtedly differentiated from the rounded neuronal cell bodies. Additionally, eGFP expression was found to be brighter in SGCs relatively to neuronal cell bodies (**Fig 2A,B**), which helps ascertain the identity of both cell types. Of note, we found that eGFP expression in SGCs and neurons reflects the endogenous S100β protein expression, showing that, in DRGs, S100β is not a glial selective promoter. These data complement studies reporting that S100β promoter drives transgene expression in some motor neurons within the brainstem and spinal cord (33, 51).

The fact that the GFAP promoter drives merely no transgene expression in SGCs from GFAP-Cre::GCaMP6f mouse line is consistent with our immunohistochemistry data showing only rare GFAP-expressing SGCs. Furthermore, our data showing Cre-mediated recombination in 58.5% of sensory neurons in such GFAP-Cre::GCaMP6f mice suggest the possibility that GFAP is expressed in DRG neuronal lineage during development. This possibility though is not supported by the absence of evidence for a developmental GFAP expression in PNS neuronal lineage, while such expression is well described in the CNS (52). Thus, together our results suggest that the GFAP promoter used in the GFAP-Cre::GCaMP6f mice is not sufficient to drive a strong and selective expression of transgenes in a large number of SGCs under physiological conditions. However, in disagreement of this hypothesis, the same promoter (2.2kB human GFAP minimum promoter, 29) has been previously used successfully to highly express a transgene selectively in the overwhelming majority of DRG SGCs with no ectopic sensory neuronal expression (53). This implies that other factors (*e.g.* number of transgene copies, gene microenvironment) may account for the variability of transgene expression levels in SGCs and cell selectivity. Indeed, transgenes insert randomly into the genome and the transgene expression cannot always mimic endogenous promoter activity. Thus, controlling gene microenvironment represents a strategy to enhance the specificity of transgenic targeting. One approach is to insert transgenes into a cassette containing all introns, promoter regulatory elements, exons and 5’ and 3’ flanking DNA of the GFAP gene (52). This approach has been used in a recent study showing specific transgene expression in SGCs of the trigeminal ganglia, although it remains to be evaluated in DRG SGCs (49). Another approach is to insert transgenes into bacterial artificial chromosomes (BACs) as they reduce the influence of chromosome position effects and allow more predictable transgene expression patterns. Despite these advantages, BAC transgenic mice appear to suffer from some remaining expression variation. Indeed, using the BAC-based GLAST-CreERT2 mice (28) to generate the GLAST-CreERT2::GCaMP6f mice, we obtained both (i) a marginal percentage of GLAST-driven CreERT2-mediated recombination in DRG SGCs, and (ii) a lack of cell specificity, while endogenous GLAST protein is selectively and prominently expressed in essentially all SGCs (47). Therefore, it stands to reason that even though both, conventional and BAC-based transgenesis, can lead to robust and cell specific transgene expression in the CNS glial cells, their expression pattern may significantly differ in DRG glia (and *vice versa*). Knock-in technology may be considered as a good alternative to traditional transgenic techniques; indeed it enables transgene insertion at a specific glial gene locus of the mouse genome, which allows excellent control of gene microenvironment, and thus is likely to better avoid chromosome position effects and circumvent the above-discussed drawbacks associated with conventional and BAC-based transgenesis. Our findings using the knock-in CX3CR1-eGFP mouse line does support the relevance of this approach by showing specific eGFP expression in the vast majority of microglial cells and macrophages in V1 and DRGs, respectively (**Fig 5**; **Fig 6**). Thus, to improve DRG glial cell specific targeting, it would be of interest in future studies to generate and/or use knock-in glial mouse models. One of such mouse models, the knock-in GLAST-CreERT2 mouse line in which CreERT2 transgene is inserted at the GLAST gene locus (54), appears to be a good candidate to examine next.

A large number of *in vivo* studies have used the GFAP or S100β promoters to drive different transgene expression in astrocytes. Results from these studies are routinely interpreted as due to the expression of transgenes only in astrocytes. Our finding that the GFAP or S100β promoters drive GCaMP6f or eGFP expression in 58.8 % or 13.5% of DRG sensory neurons, respectively, should be considered when interpreting *in vivo* results from such studies.

In conclusion, most of the tools tested in the current study were found ineffective in studying selectively the majority of SGCs in DRGs, although a lot of molecular and functional similarities exist between DRG SGCs and astrocytes. Therefore, further work is required for characterizing and identifying other already available tools as well as developing new genetically-modified mouse lines and adeno-associated viral tools to specifically target large proportions of DRG SGCs, but also macrophages. Together with the two mouse lines validated here (Cx43-CreERT2::GCaMP6f and CX3CR1-eGFP), these future molecular tools will be of prominent interest in understanding better how DRG glia can modulate sensory information processing under physiological and pathological conditions.

## Acknowledgments

We gratefully acknowledge S. Antoine and S. Guinoiseau for animal care; S. Guinoiseau and P. Meriau for dissecting some tissues; C. Ayissi Sama and M. Tantouch for slicing some tissues; F. Charbonnier, B. Delhomme, P. Djian, and M. Oheim for sharing pieces of equipment (cryostat, microscopes); J. Strinnakre, S. Guinoiseau and P. Meriau for proof reading; and both the imaging and mouse core facilities, which are supported and funded by the CNRS, INSERM and Paris Descartes University.

## Author contributions

YR, BR, EM and CA designed and interpreted experiments. YR wrote an initial draft of the manuscript and CA edited and wrote the manuscript. YR, EM, and BR performed experiments, data analysis and wrote the legends. CA conceived and supervised the project.

## Supporting Online Material

**Supplementary Figure 1.**
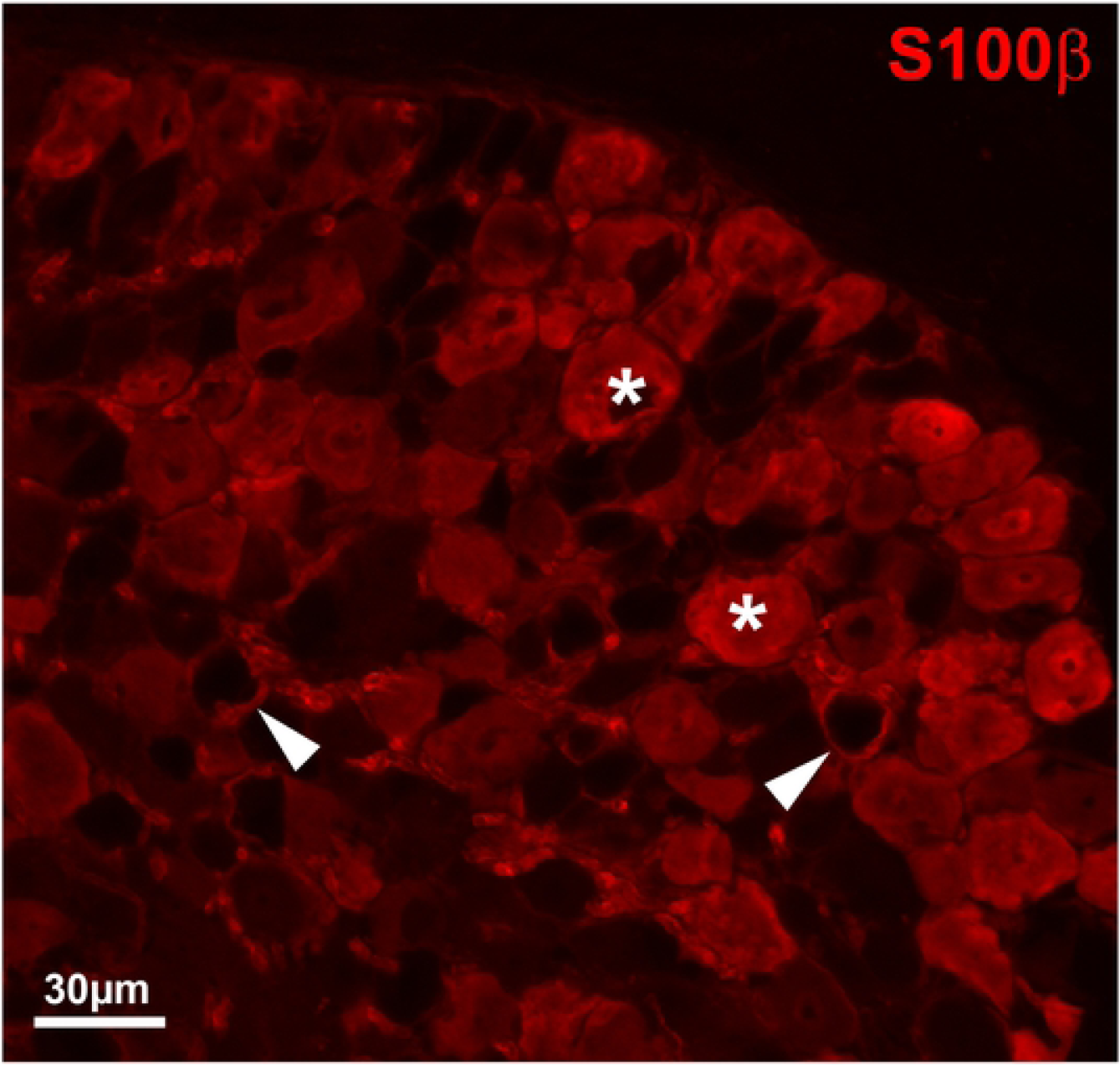
S100**β** immunoreactivity signal (red) in DRGs from wildtype mice, showing that endogenous SIOOP is expressed in both ring-shaped SGCs (arrowheads) and sensory neuron cell bodies (asterisks).

